# Tracing spontaneously occurring mutations in *Fusarium graminearum* laboratory strains resulting in reduced virulence on wheat

**DOI:** 10.1101/2025.08.06.668883

**Authors:** Thomas Svoboda, Florian Kastner, Michael Freitag, Joseph Strauss

## Abstract

**Background:** *Fusarium graminearum* is a well characterized plant pathogenic fungus which is able to infect a broad range of economically relevant crop plants. Besides yield reduction this fungus is also responsible for mycotoxin contamination of food and feed. Upon propagation under laboratory conditions, mutations may occur which would be disadvantageous for fungal fitness in nature but not in the lab where the strains are usually grown on nutrient-rich media under optimal growth conditions. In this study we characterized four phenotypically different *Fusarium graminearum* strains for fitness traits and compared their genomes to trace down mutations responsible for the phenotypes.

**Results:** The four tested *F. graminearum* PH1-derived strains revealed differences in their phenotypic appearance and also in their secondary metabolite profiles expressed on different growth media. Also, two of the investigated strains (94 and 96) showed significantly reduced virulence on wheat upon point inoculation of flowering wheat. We identified one high impact mutation in each of the two strains. In strain 96 a loss of function mutation occurred in FGSG_00355 which has a high similarity to Ras GTPase activating proteins and consequently may have an impact on the cell cycle. Even though strain 96 showed enhanced DON production *in vitro*, the strain was no longer able to spread within the wheat ear in infection assays. In strain 94 we identified an insertion of an A rather at the end of FGSG_00052 leading to a frameshift and consequently mutation of the last three amino acids and a shift of the stop codon by seven amino acids. Even though knock-out of this putative transcription factor has been described by Son et al (2011) to have no impact on virulence, changes at the C-terminal region may result in changes of the binding affinity.

**Conclusions:** We tracked down the mutations which might be responsible for the changes in phenotypic appearance, secondary metabolite profile as well as virulence. Yet, a closer biological characterization is necessary to determine the impact of these mutations on the fungus.

**Impact statement:** Mutations under laboratory conditions can occur spontaneously. Due to the lack of selection pressure, also potentially deleterious mutations remain unnoticed as long as all nutrients are provided. In this study we analyzed four phenotypically different *Fusarium graminearum* PH-1 strains among which two showed significantly reduced virulence on wheat. In one strain we identified a loss of function mutation in a Ras-GTPase activating protein resulting in enhanced growth on complete media but at the same time strongly reduced virulence. The mutation in the second strain was an insertion in the C-terminal region of a transcription factor. The exact role of the Ras-GTPase activating protein during infection and the impact C-terminal elongation of a yet uncharacterized transcription factor is yet to be investigated. This study underscores the importance of regularly checking laboratory strains on their traits that such mutations which may have an impact on your research data, do not remain unnoticed.

**Data summary:** The code used for web scraping is available on github (https://github.com/cicci726/webscraping/tree/main). The sequencing files are deposited at NCBI in the sequence read archive (BioProject ID: PRJNA1293145).

## Introduction

Mutations in fungi occur through various mechanisms and at different rates. Depending on the localization the impact of a mutation can either be neutral, deleterious or beneficial. In general, beneficial mutations will establish in nature while under laboratory conditions also deleterious mutations will persist. The **<u>R</u>**epeat-**I**nduced **<u>P</u>**oint (RIP) mutation pathway which is specific to certain fungal taxa, targets repeated DNA sequences and can drive genome divergence and chromosome architecture changes. Several studies demonstrated that RIP is another tool for heterochromatic gene silencing of repetitive DNA thereby limiting accumulation of transposable elements (Freitag et al. 2002; Bouhouche et al. 2004; Hood, Katawczik, and Giraud 2005; Lewis et al. 2009; Gladyshev 2017). The same system was observed subsequently also in other genera including *Fusarium circinatum* (van Wyk et al. 2019) and other Ascomycota (van Wyk et al. 2020). Some fungi, such as the *Schizophyllum commune*, accumulate mutations rapidly, at a rate of 1.4×10⁻⁷ substitutions per nucleotide per meter of growth (Bezmenova et al. 2020). The RIP process, which occurs during the sexual reproductive cycle, can both enhance and impede genetic diversity in fungi. It affects not only repeated genomic regions but also adjacent non-duplicated genes, potentially driving rapid adaptation in some species (Hane et al. 2015).

In *Fusarium graminearum*, a major cereal pathogen, mutations occur at varying frequencies influenced by environmental factors. Genome sequencing revealed low repetitive sequences and paralogous genes, possibly due to repeat-induced point mutations (Cuomo et al. 2007). Highly polymorphic regions containing plant-fungus interaction genes were identified, suggesting adaptive evolution. A consecutive analysis of the genomic sequence corrected assembly errors and also included AT rich sequences, centromeric and subtelomeric regions as well as the telomers (King et al. 2015). Research on soil fungi in “Evolution Canyon” demonstrated that strains from more stressful environments exhibited higher spontaneous mutation frequencies compared to those from less stressful areas, even when grown under mild laboratory conditions (Lamb et al. 2008). This suggests inherited differences in mutation rates as a potential adaptive feature.

*F. graminearum* exhibits diverse phenotypic traits. A multivariate analysis revealed that *F. graminearum* and *F. meridionale* strains are structured by species, with distinct characteristics in growth, reproduction, and pathogenicity (Machado et al. 2023). These phenotypic differences can be taken into account for disease management considering tebuconazole resistance. Adaptation of *F. graminearum* to the fungicide tebuconazole has been demonstrated to result in two distinct phenotypes with varying levels of fitness, fungicide resistance, virulence, and mycotoxin production (Becher et al., 2010).

Laboratory strains are usually cultivated under ideal conditions which leads to the absence of main dogma in nature regarding “survival of the fittest” (Darwin and Kebler 1859). This lack of selection pressure can result in mutations which are advantageous but also deleterious to the fungal fitness. The term “degeneration” has been initially defined by Reusser (1963) where the changes of microbial cultures upon repeated transfer were described. For example, serial subculturing of *Aspergillus fumigatus* on plates has been demonstrated to cause alterations regarding virulence and sensitivity to antifungal agents (Curtis, Walshe, and Kavanagh 2023). Also, the degeneration of biotechnologically relevant fungal species is often detrimental to the yield as described for biotechnological processes with *Aspergillus niger*, *Aspergillus oryzae*, *Trichoderma reesei*, and *Penicillium chrysogenum* (Danner, Mach, and Mach-Aigner 2023).

In this study we characterize four *F. graminearum* PH-1 strains with different phenotypic appearances on solid media for their virulence phenotypes to test if there is a correlation between morphology and virulence. Additionally, to the classical methods for phenotypic description, we use a comparative genomics approach to find mutations which may be responsible for the reduced virulence of two of the characterized strains.

## Material and Methods

### Strains

In this study four different *Fusarium graminearum*, PH-1, wild type strains were used. They originated from the laboratories of Gerhard Adam (93), Michael Freitag (94), Jens Sørensen (95) and Joseph Strauss (96).

### Sporulation and growth conditions

Sporulation was performed by inoculation of agar plugs in 50 ml mung bean broth followed by incubating at 20°C, 140 rpm for 3 days. The spores were harvested by filtration through glass wool filters, followed by centrifugation and counting. The growth tests were performed on Fusarium minimal media (FMM: KH_2_PO_4_ 1g/L, MgSO_4_*7H_2_O 0.5 g/L, KCl 0.5 g/L, Sucrose 30 g/L, NaNO_3_ 2 g/L, Trace elements 0.2 ml (100 mL trace elements stock: Citric acid 5 g, ZnSO_4_ * 6 H_2_O 5 g, Fe(NH_4_)_2_(SO_4_)2 * 6 H_2_O 1 g, CuSO_4_ * 5 H_2_O 250 mg, MnSO_4_ 50 mg, H_3_BO_4_ 50 mg, Na_2_MoO_4_ * 2 H_2_O 50 mg)) and on Fusarium complete media (FCM: Glucose 10 g/L, Meat peptone 2 g/L, Casamino acids 1 g/L, Yeast extract 1 g/L, ASPI-salts 50 mL/L (Aspi salts stock: KCl 10,4 g/L, KH_2_PO_4_ 16,3 g/L, K_2_HPO_4_ 20,9 g/L, Mg(SO_4_)_2_ 100g/L), Mg(SO_4_)_2_ 5 mL/L, Trace elements 1 mL/L (Trace elements stock solutions (80 ml): Sol. A: EDTA 10g FeSO_4_*7H_2_O 1g; Sol. B: ZnSO_4_*7H_2_O 4,4g, H_2_BO_3_ 2,2g, MnCl_2_*4H_2_O 1g, CoCl_2_*6H_2_O 0,32g, CuSO_4_*6H_2_O 0,32g, (NH_4_)6Mo_7_O_24_*4H_2_O 0,22g)) in triplicate. A spore suspension of 10^5^ spores/ml was used from which 10 µl were pipetted onto the plate. The plates were incubated at 20°C for five days.

### Wheat infection

For wheat infection, all strains were sporulated as described above. The spores were counted and diluted to a concentration of 4*10^5^ spores/ml. Two spikelets in the middle of the ear were infected using 10 µl of the spore suspension per floret. The wheat heads were covered with plastic bags which were previously sprayed on the inside with sterile water to maintain humidity during infection. After two days the bags were removed and the progress of infection was visually determined every two days.

### Toxin tests

The toxin tests were conducted on rice. Therefore 2 g rice were soaked with 2 ml water for 1 hour and autoclaved. 1*10^5^ spores of the respective fungus were put on the rice media and incubated at 20°C in the dark for 2 weeks. Metabolites were extracted using ACN:H_2_O:HAc = 79:20:1, incubation for 1 hour at 20°C, 180 rpm followed my LC-MS analysis.

### Bioinformatic analysis

Once the sequencing was completed, the quality was checked using fastQC v0.11.9. To remove the sequencing primers trimmomatic v0.39 was used. The Burrows-Wheeler aligner (Li 2013) was used to align the Illumina reads to the reference genome. Freebayes v1.3.2 was used for variant calling (Garrison and Marth 2012). The contigs were re-named according to the reference available in snpEff (Cingolani et al. 2012). snpEff analysis for the respective mutations was done in R.

To get information on the predicted function of the mutated genes, web scraping was done. Therefore, we used the information available on the homepage of the National Center for Biotechnology Information (NCBI). The corresponding python notebook for web scraping is available on github (https://github.com/cicci726/webscraping/tree/main).

## Results and discussion

We were emanating from four *Fusarium graminearum* PH-1 wild type strains which all showed different macroscopic morphologies FMM and FCM solid growth medium (Figure 1A). While strain 93 shows the normal radial growth on FMM and FCM, strain 96 forms abundant mycelia on FCM while on FMM only faint growth is observed. Strain 95 appears yellow on FCM while on FMM a red pigment is produced. *Fusarium* strain 94 has a phenotypic appearance on FCM comparable to strain 93 while on FMM mycelium formation is reduced. A comparison of the spores produced by the respective strain revealed that the spores formed by strain 96 are less curved (Figure 1B) while all other strains showed no deviation from the known banana shape spore morphology.

**Figure 1:**
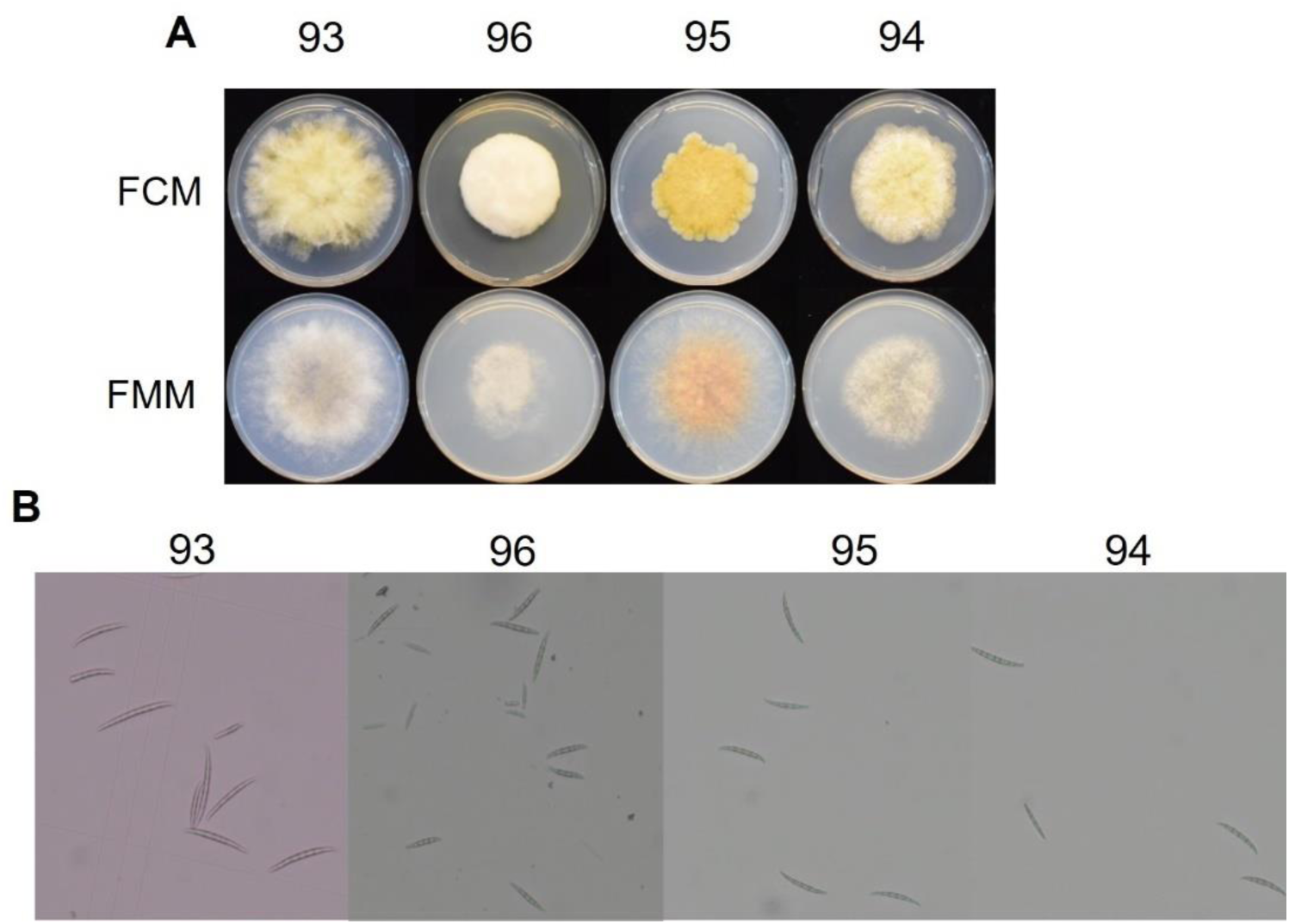
Characterization of the four phenotypically different *F. graminearum*, PH-1, wild type strains; A) Phenotypic appearance on FMM and FCM, B) spores of the different strains under the microscope, C) Selected secondary metabolites produced by the two *Fusarium graminearum* PH-1 exhibiting lower virulence on wheat compared to a healthy wild type strain.

### Two of the tested F. graminearum wild type strains are hardly able to spread within the wheat ear after point inoculation

To find out whether the different phenotypes also show differences in virulence a wheat infection was performed. The point inoculation of the susceptible wheat cultivar Apogee revealed that strain 94 and strain 96 are able to infect the inoculated spikelets, however, both strains are hardly able to spread within the ear. Twelve days post inoculation these two strains hardly go beyond the two inoculated spikelets (Figure 2A). While strain 94 and 96 do not significantly differ from each other over time, there is a significant difference between these two strains and the other two, 93 and 95, after 10 dpi (Supplementary Table 1).

**Figure 2:**
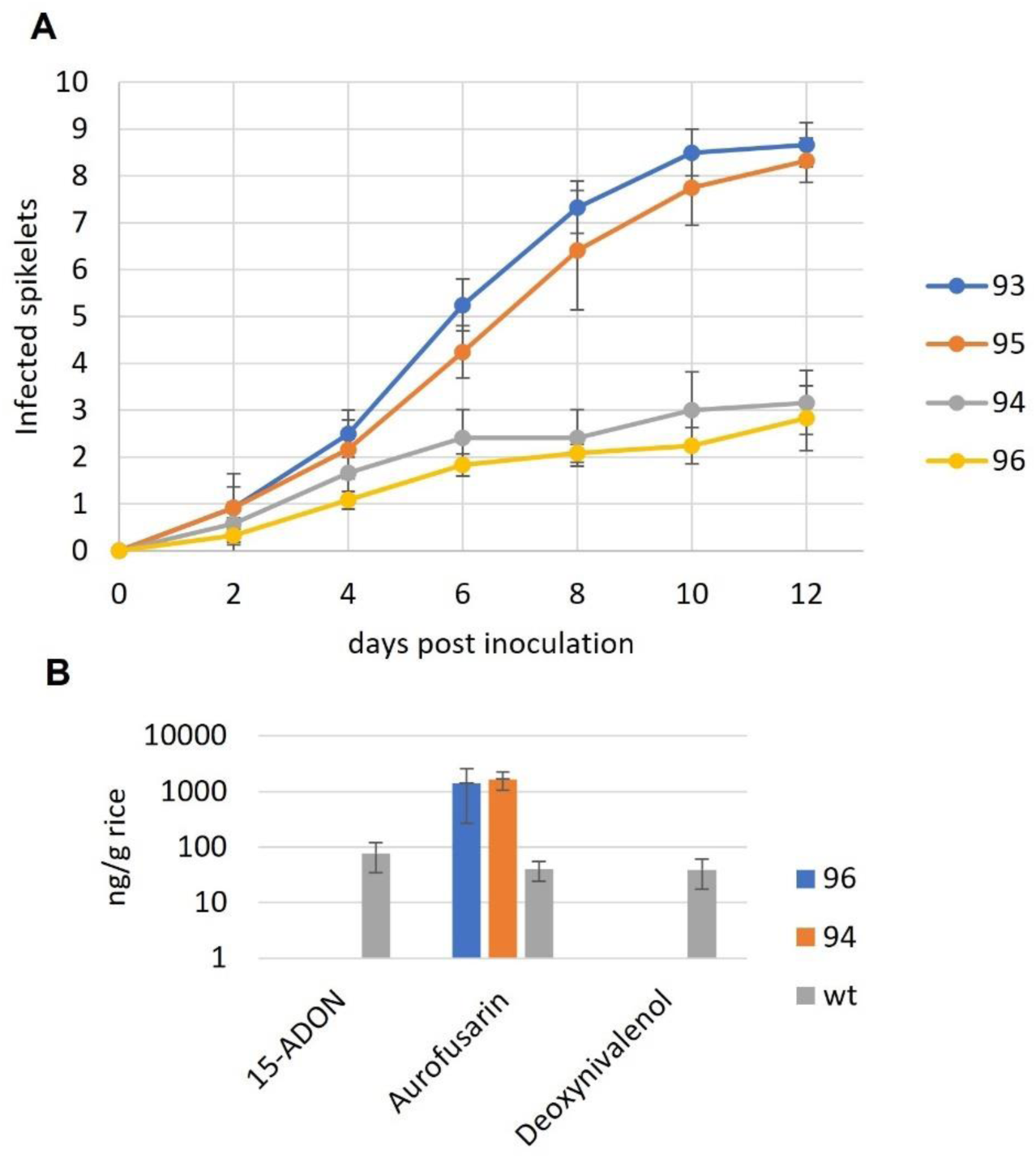
A) Virulence test on wheat (cv. Apogee), Secondary metabolite analysis of the two strains which show reduced virulence compared to a PH-1 wild type strain.

Further tests of the strains revealed that there were also differences in secondary metabolite production. As shown in Figure 2B, both of the strains which show a significantly reduced virulence on wheat produce significantly lower amounts of DON on rice media. In contrast, aurofusarin production was significantly higher in strain 94 compared to the wild type.

### Genome comparison

To find out whether genomic alterations are responsible for the different phenotypic appearances and differences in virulence the genomes were sequenced. All Illumina reads were aligned to the published reference genome of *Fusarium graminearum*, PH-1 (https://fungi.ensembl.org/Fusarium_graminearum/Info/Index). A first comparison of all mutant strains revealed that under the parameters used for variant detection 3006 mutations were shared between all of the strains (Fig. 3). Since the sequencing results were consistent and the read depth was high and unambiguous in all strains, it is highly likely that these nucleotides were not correctly assigned in the reference genome. Apart from the mutations occurring in all strains each strain has a number of unique mutations. Besides, also mutations occurring in two or three of the strains were identified. The results of the phenotypic characterization directed our main interest to the two strains showing reduced virulence on wheat. While in strain 94 and strain 96 109 and 156 unique mutations were detected, respectively, these strains also exclusively share 21 more mutations which were not identified in the strains 93 and 95 (Fig. 3).

**Figure 3:**
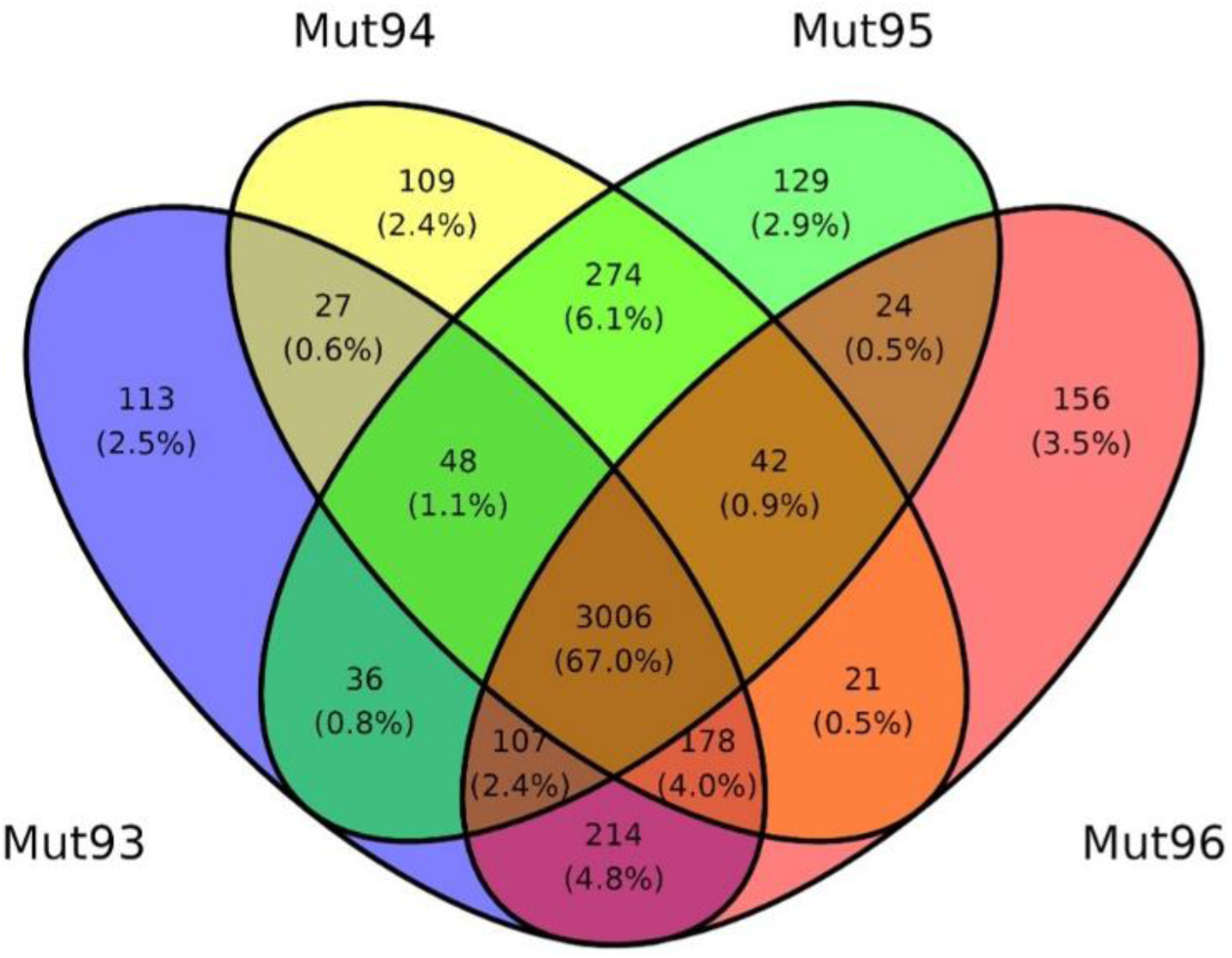
Venn diagram of the mutations detected in the different strains.

Assessment of the impact of the different mutations revealed that among the mutations shared between the strains 94 and 96 only one moderate impact in FGSG_01434 was identified. It is annotated as complex mutation including several deleted bases and a few mutations at the region. FGSG_01434 is a yet uncharacterized protein belonging the amino acid/polyamine/organocation (APC) superfamily. However, since also different mutations have been detected in that gene in the other strains, we assume that the impact of this mutation is negligible.

In contrast, strain 94 showed 109 unique mutations among which also high impact effects were predicted. A closer verification of the aligned sequences revealed that there is only one gene, FGSG_00052, where no other mutations were detected in the other three strains (Supplementary Figure 1). This high impact mutation exclusively present in strain 94 may have a considerable impact on virulence. In FGSG_00052 we identified one insertion of an adenine at position 942 leading to a frameshift three amino acids in front of the stop codon. The resulting protein is seven amino acids longer compared to the native one and the last ten amino acids are deviating. While FGSG_00052 is annotated as hypothetical protein on NCBI, the same protein is described as Zn(II)Cys6 transcriptional activator on fungidb. A structural analysis of the impact of the mutation on the structure with AlphaFold 3 (https://golgi.sandbox.google.com/) revealed that the C-terminal region is located on the outside of the protein (Supplementary Figure 2).

While in the native protein the last six amino acids are RKKVII which are either positively charged or non-polar, the mutated protein shows RKKSHHLSLLKNW. We can see that most of the mutated/ added amino acids have the same properties as the ones in the native protein there are also two serins introduced which are polar and may interfere with the integrity of the C-terminal region. The C-terminal region of transcription factors often plays a specific role. In the mitochondrial transcription factors Mtf1 and TFB2M of *Saccharomyces cerevisiae* and *Homo sapiens*, respectively, the C-terminal region has been shown to be crucial for DNA binding (Basu et al. 2020). Alternatively, the C-terminal domain can also play a role in transactivation (Stracke, Turgut-Kara, and Weisshaar 2017; Valsecchi et al. 2013; Wisniewski et al. 1996). However, in FGSG_00052 the DNA binding domain seems to be located at the N-terminal region (Supplementary Figure 3) while the C-terminal has no closer predicted function apart from fungal transcription factor. Data of Zhao et al. (2014) indicate that in the mycelium FGSG_00052 is hardly expressed (TPM = 0.01) while in spores the expression of this transcription factor is at 1.32 TPM.

In all aspects of the fungal life cycle, transcription factors are crucial regulators of gene expression. In this context, a comprehensive study of 657 transcription factor mutants in *F. graminearum* provided insights into the regulation of various traits, including growth, development, and toxin production (Son et al. 2011). This deletion mutant library has been used to assess the impact the respective transcription factor on *Fusarium graminearum* virus 1 (FgV1) infection, revealing associations between viral RNA accumulation and phenotypic differences in infected mutants (Yu and Kim 2021). These studies collectively demonstrate the complex phenotypic diversity of *F. graminearum* and its adaptability to different environmental pressures. Among the putative transcription factors disrupted by Son et al. (2011) FGSG_00052 was included. The characterization of the phenotype showed comparable virulence on wheat, however, the perithecia formation was reduced by 75-100% compared to the wild type strain (Son et al. 2011).

Yet, the exact role of this putative transcription factor has not been described. It seems to be involved into perithecia formation, however, it will most likely play a role in other pathways as well. Even though the knock-out mutant shows comparable virulence to the wild type (Son et al. 2011) it remains to be elucidated whether the identified mutation has an impact on potential stronger activation or repression of different pathways leading to changes in the secondary metabolite profile as well as reduced virulence.

Among the unique mutations present in strain 96, there is one mutation where a high impact is predicted in gene FGSG_00355, annotated as GTPase activating protein (Supplementary Figure 4). The mutation in this gene is at amino acid 72 where a C ⟶ T mutation was detected. Hence the codon CGA which encodes for arginine was mutated into TGA, a stop codon, resulting in a loss of function. The results of a blastp search showed that FGSG_00355 has 92.02% of the amino acids identical with the characterized Ras-GAP of *Colletotrichum obiculate* MAFF 240422 (Supplementary Figure 5) (Harata and Kubo 2014). The activity of Ras is regulated in a cycle where the guanine exchange factor (GEF) exchanges GDP for GTP and the dephosphorylation of Ras-GTP (Hennig et al. 2015). Ras itself already has an intrinsic GTPase activity, however, the activity is strongly enhanced by interacting with the Ras-GTPase activating protein (Schaber et al. 1989). Hence a loss of function mutation in Ras-GAP will result in uncontrolled growth due to the presence of the constantly active Ras-GTP. In *Saccharomyces cerevisiae* the function of IRA2, a homolog of the human Ras-GAP, has been reported to be crucial to stimulate the GTPase activity of Ras (Tanaka et al. 1991). In *Aspergillus nidulans*, a Ras-GAP mutation has been shown to be responsible for compact, fluffy growth. Knock-out of gapA also results in reduced conidiation, however, giant spores with multiple nuclei are formed (Harispe et al. 2008). In *Fusarium graminearum*, the disruption of Fgcdc25, which was supposed to be the only Ras guanine nucleotide exchange factor (RasGEF), resulted in total loss of pathogenicity *in planta* (A. Chen et al. 2020). Also, toxisome formation and DON production were reduced in the mutant strain (A. Chen et al. 2020). In contrast, our virulence test showed that the mutant is at least able to infect to inoculated spikelets. While Chen et al. (2020) reported that mutation of Fgcdc25 results in decreased DON production, the loss of function mutation of FGSG_00355, a putative Ras-GAP, resulted in increased DON production in axenic culture. This is plausible since the mutation of the gene with the opposite function causes the opposite effect. However, one would assume hypervirulence upon constant activation of Ras which is not the case. Rather only the inoculated spikelets can be infected but spreading of the fungus within the ear is hardly possible. Yet, mutation of the highly similar *CoIRA1* gene in *Colletotrichum obiculare* also resulted in significantly reduced but not completely abolished virulence (Harata and Kubo 2014).

Two Ras GTPases in *Fusarium graminearum* have been described (Bluhm et al. 2007). While Ras1 is essential for the fungus, knock-out of Ras2 resulted in reduced perithecia formation and decreased virulence on wheat while the DON levels were comparable to the wild type (Bluhm et al. 2007). In *Colletotrichum obiculare* CoIra1, which has a high similarity to FGSG_00355, has been described to regulate the intracellular cAMP levels through CoRas2 (Harata and Kubo 2014) similar to Ira1 and Ira2 in *Saccharomyces cerevisiae* which act through Ras1 and Ras2 (Harashima et al. 2006). cAMP signaling plays a crucial role in the regulation of pathogenicity and conidial germination (Takano et al. 2001) which is overlapping with our phenotypic characterization of strain 96.

### Conclusion

In this study, naturally occurring mutations were investigated using a comparative genomics approach. We identified mutations in the two strains which show a reduced virulence on wheat. Due to the phenotype and the description in literature, there is a high probability that the mutated FGSG_00355 is indeed a Ras-GTPase activating protein. Yet, it is not clear why this strain shows a lower virulence on wheat even though it shows strong growth on plates. Regarding the mutation in strain 94, a putative transcription factor, only hypotheses can be formed. However, it seems to be obvious that this gene might have a direct or indirect impact on in the regulation of secondary metabolism. For both mutations identified in this study an experimental investigation is required to find out which alterations are caused by these mutations resulting in reduced virulence. If confirmed, investigation of the impact of the mutations on transcriptomic level during infection may give us a deeper insight into the role of the respective gene.

## Contributions

TS: draft manuscript preparation, experimental design; phenotypic characterization, bioinformatic evaluation; FK: phenotypic characterization; MF: data evaluation; JS: experimental design, data evaluation, funding acquisition.

## Acknowledgements

The authors thank Anna Atanasoff-Kardjalieff for the preparation of preliminary data as well as Gerhard Adam and Jens Sørensen for providing *F. graminearum* wild type strains

---

**Supplementary Figure 1:**
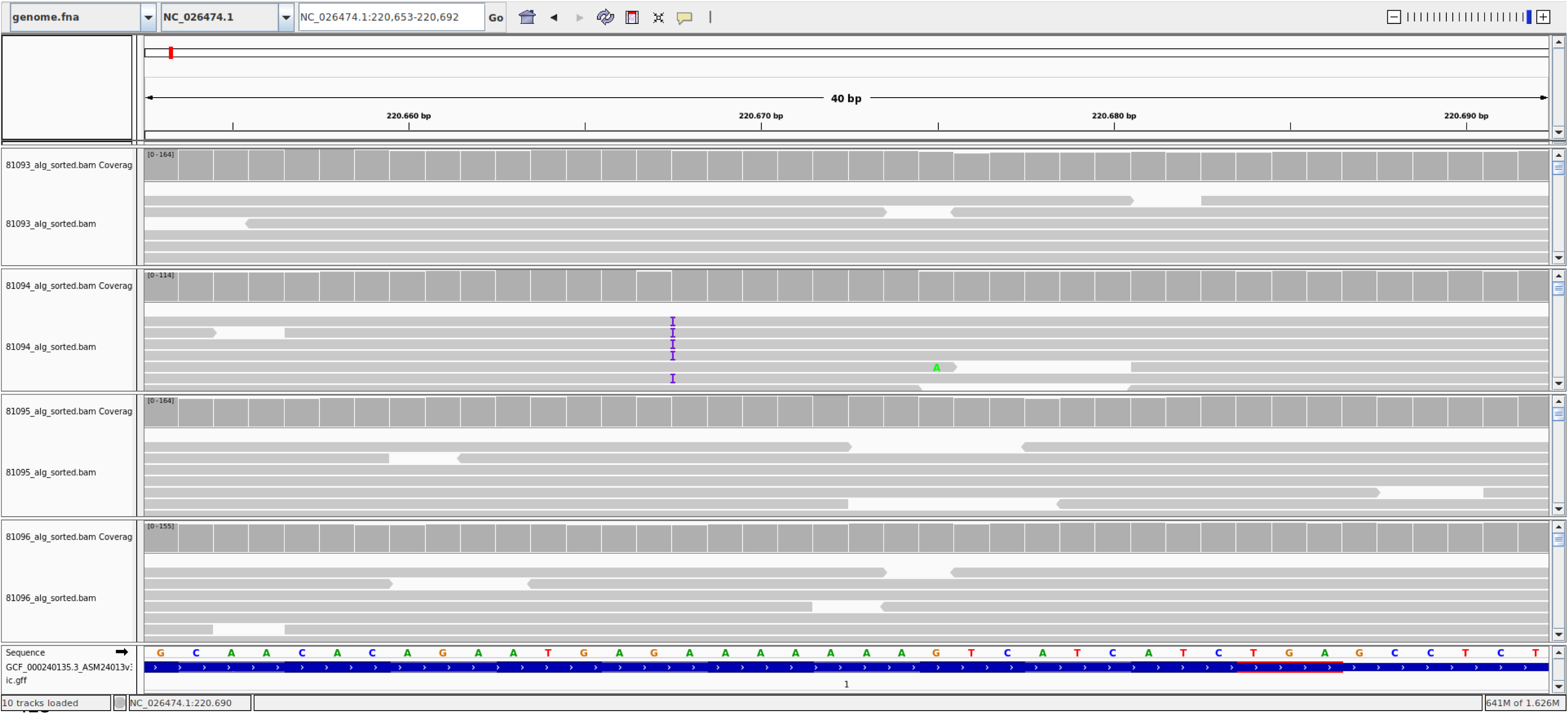
Unique high impact mutation in strain 94 in FGSG_00052.

**Supplementary Figure 2:**
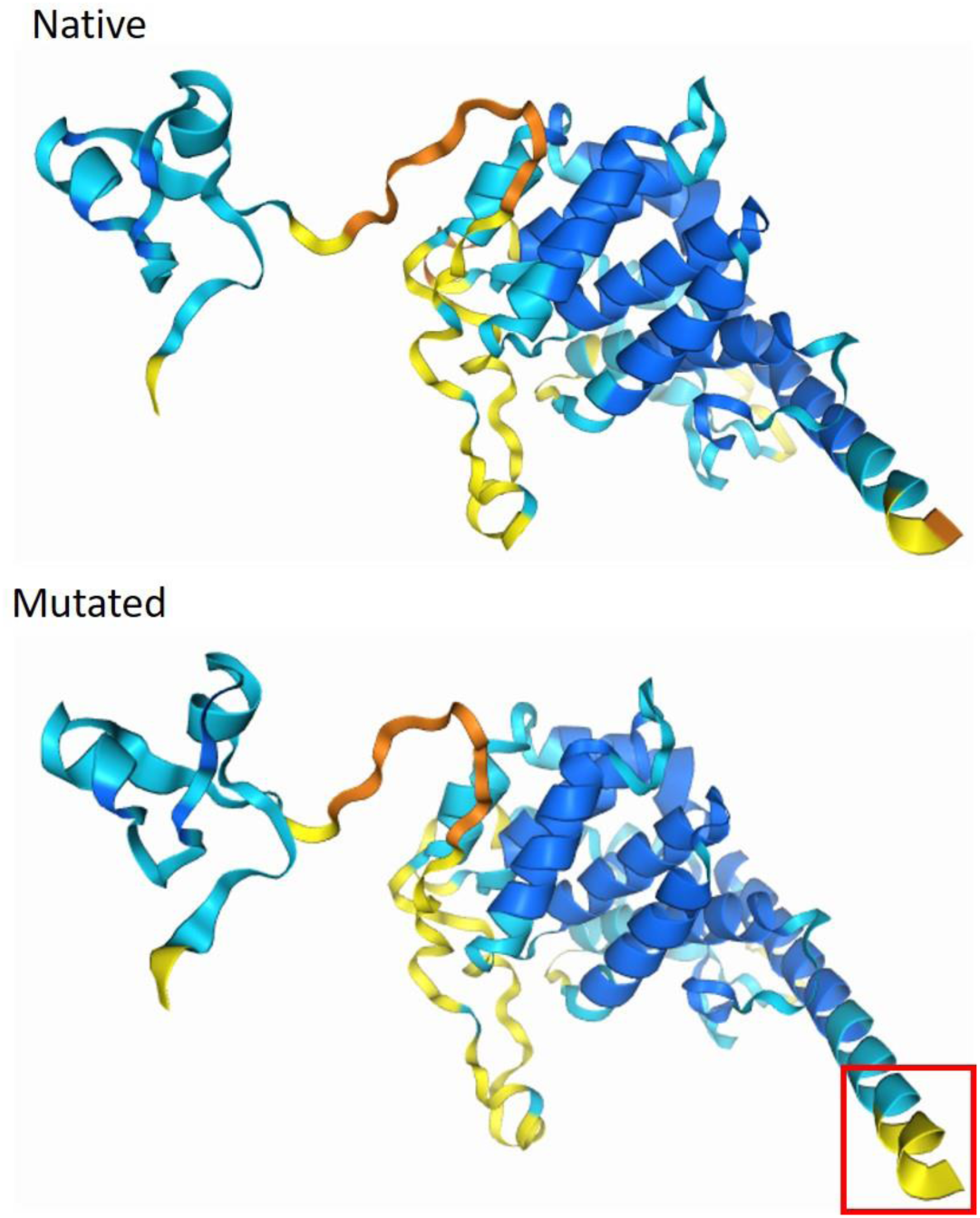
Structural analysis of the native and the mutated version of FGSG_00052 using AlphaFold. The C-terminal end of the mutated protein is 7 amino acids longer compared to the native one (red rectangle).

**Supplementary Figure 3:**
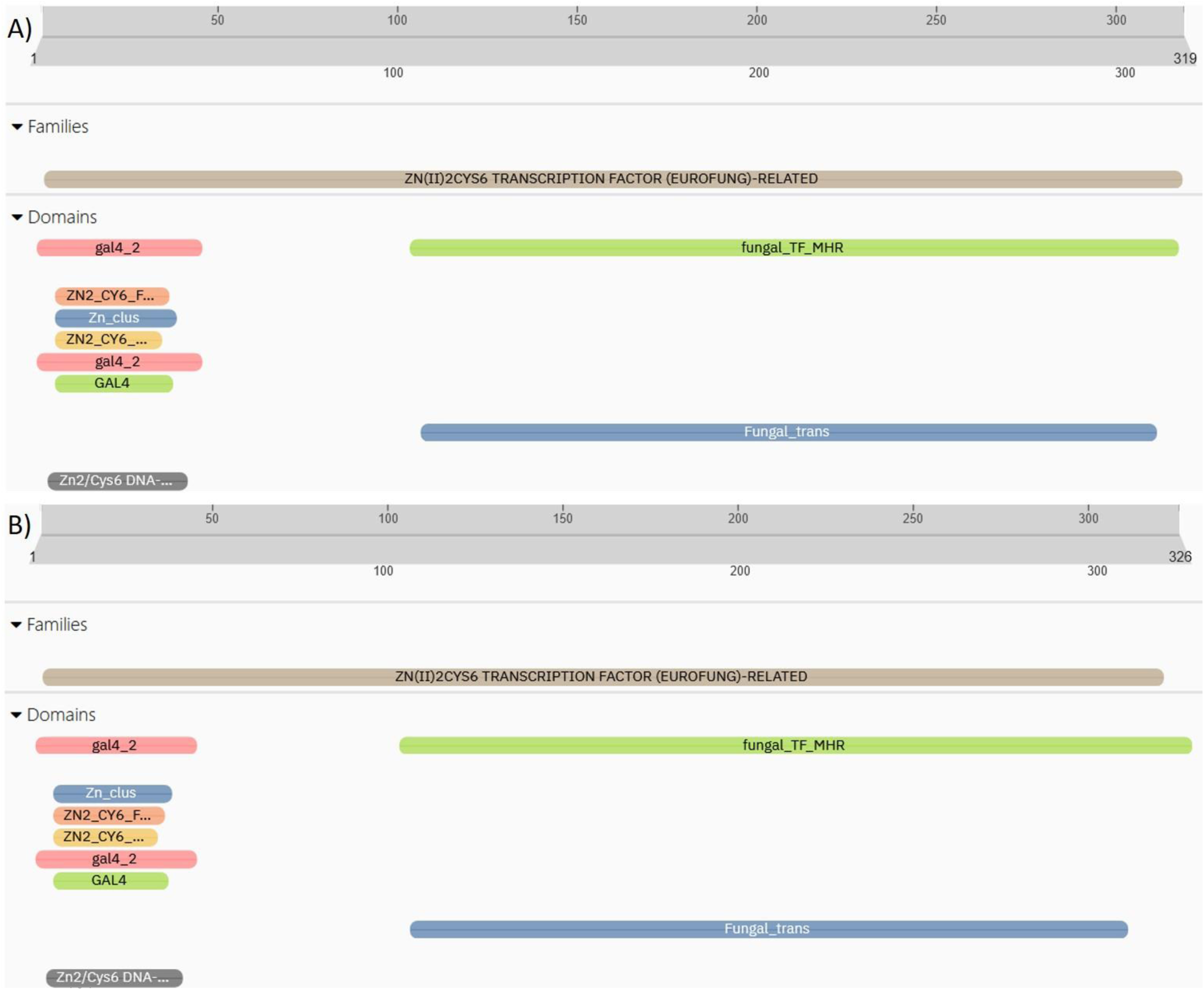
Predicted conserved domains of the native FGSG_00052 (A) and the mutated version (B) (https://www.ebi.ac.uk/interpro/); at the N-terminal region the DNA-binding domain is located. The predicted fungal transcription factor domain (Fungal_trans) is stretching between amino acids 108-308 in both proteins.

**Supplementary Figure 4:**
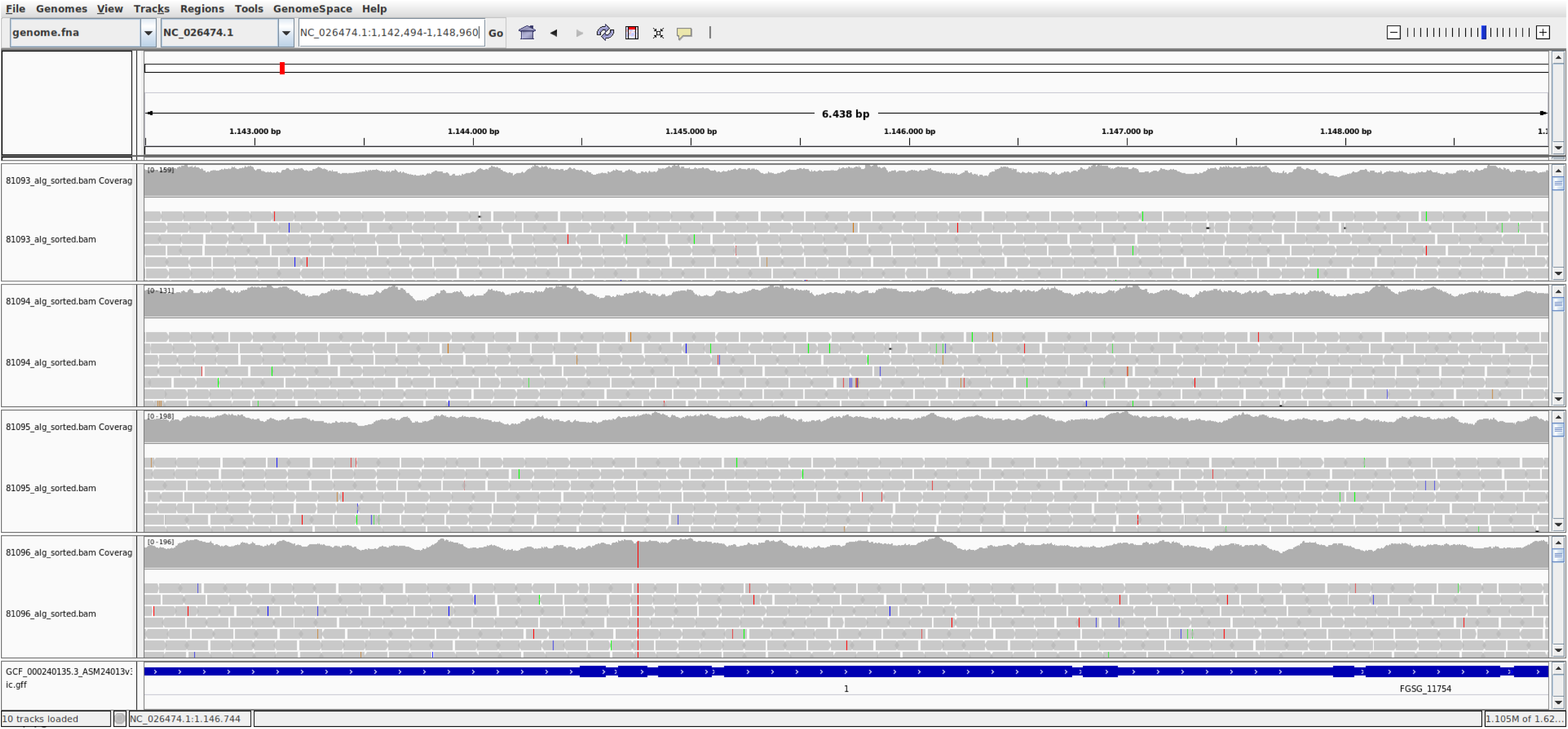
Unique high impact mutation in strain 96 in FGSG_00355.

**Supplementary Figure 5:**
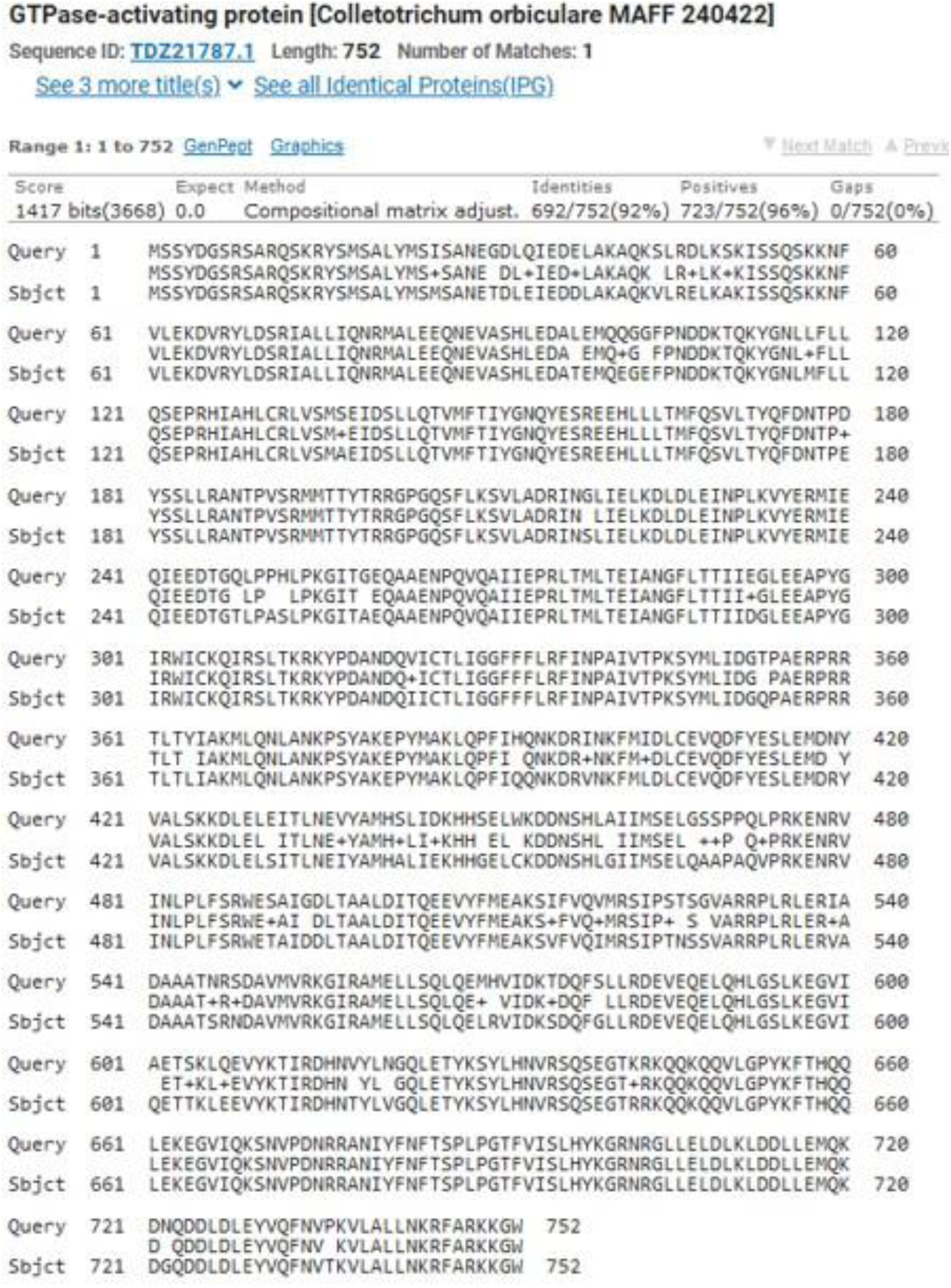
Alignment of FGSG_00355 to CoIra.

