## Supplementary figures and images for "Tracing spontaneously occurring mutations in *Fusarium graminearum* laboratory strains resulting in reduced virulence on wheat"

### Supplementary Figure 1

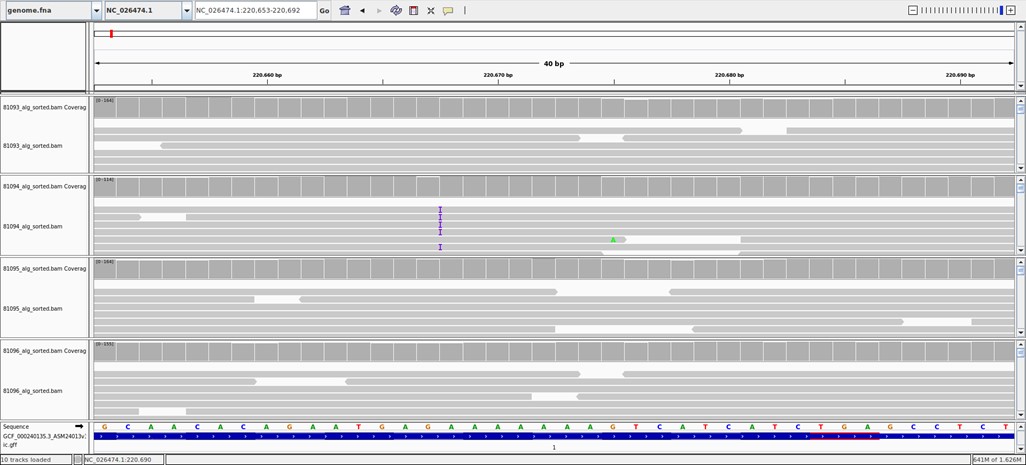

### Supplementary Figure 2

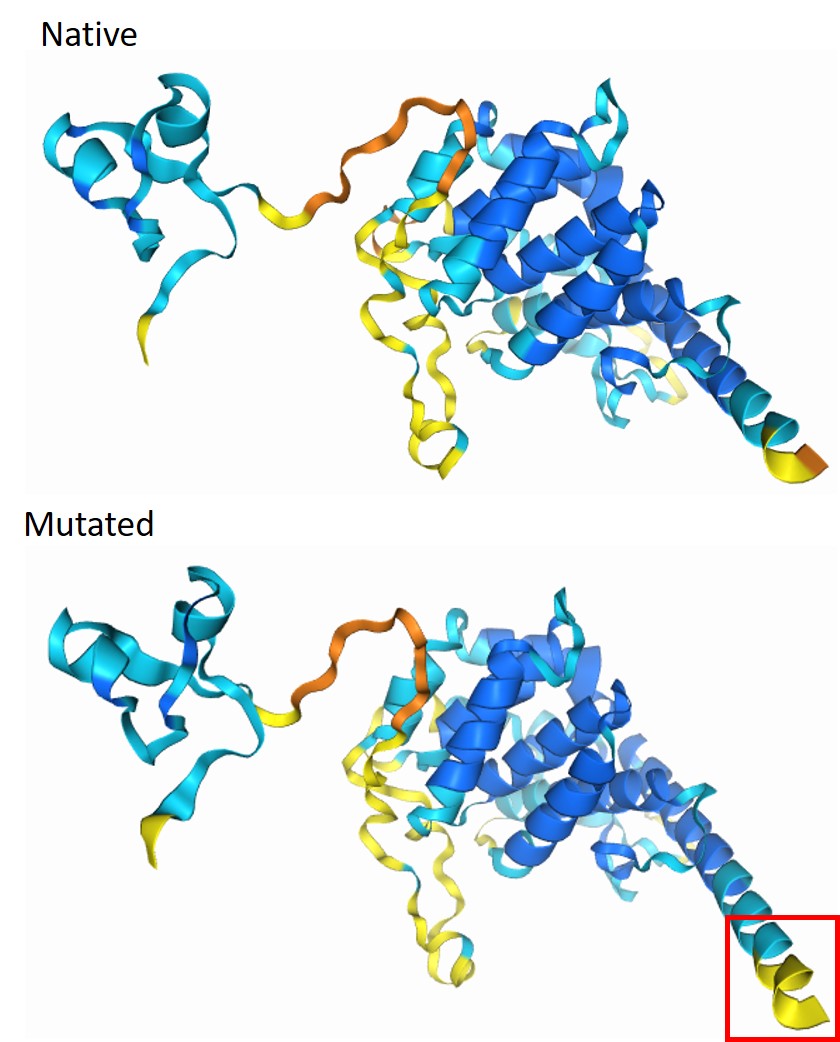

### Supplementary Figure 3

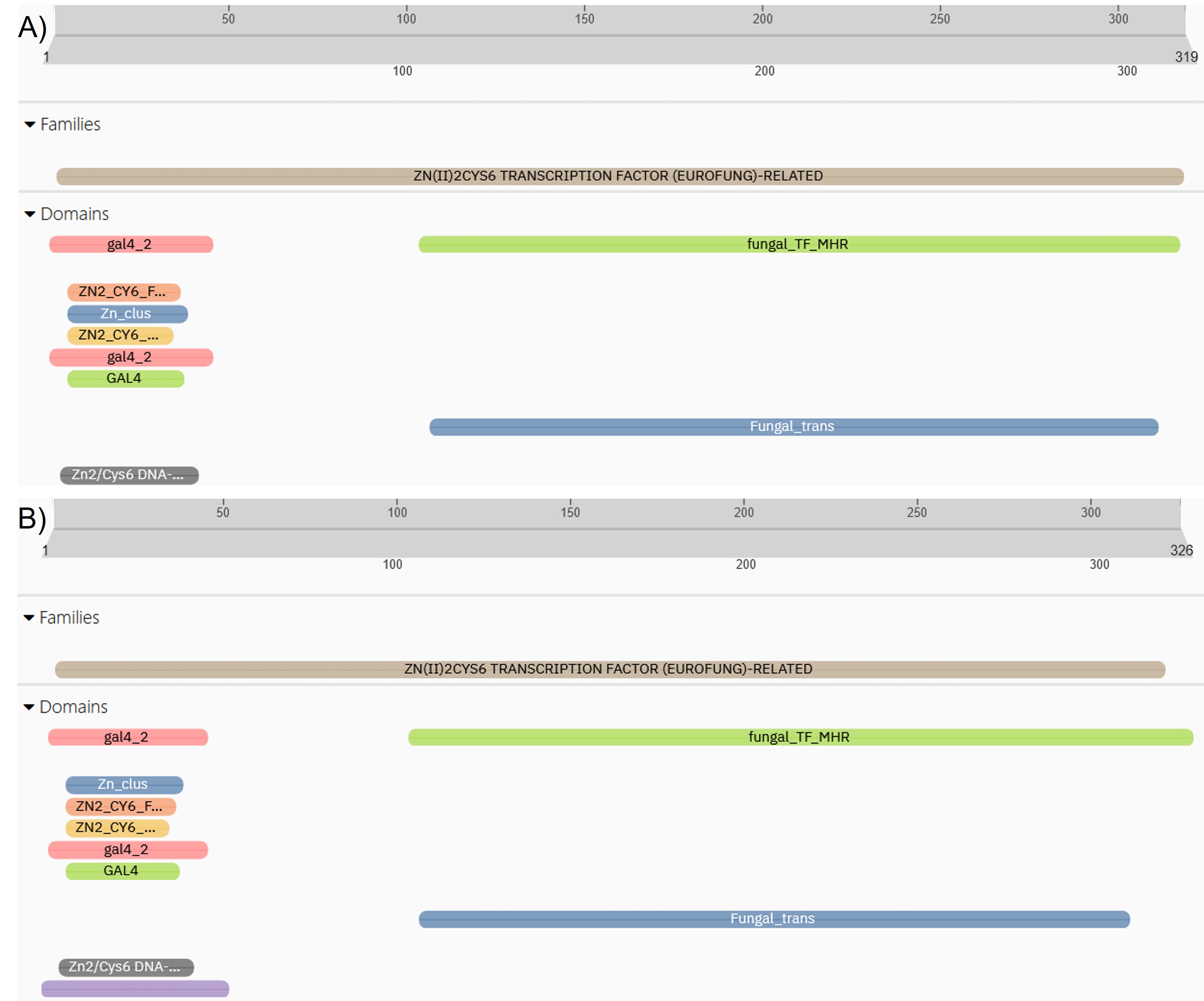

### Supplementary Figure 4

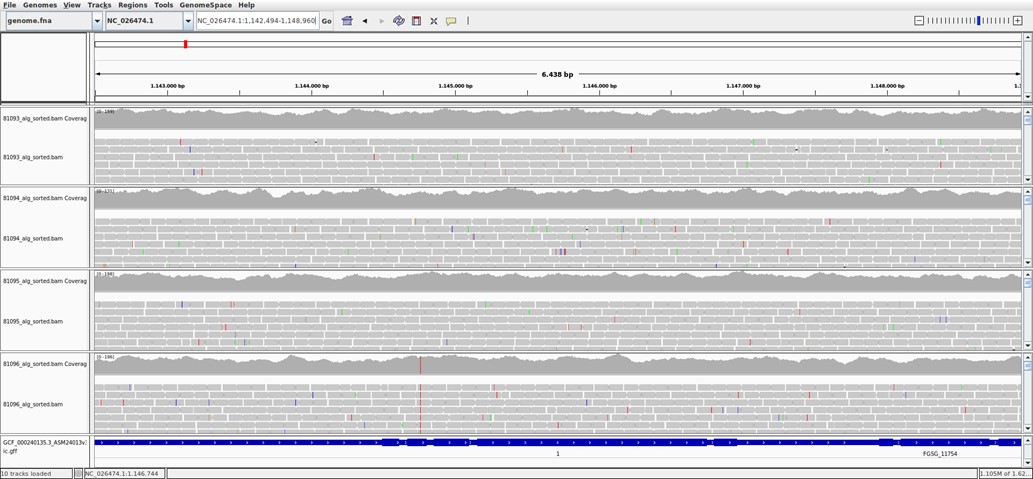

### Supplementary Figure 5

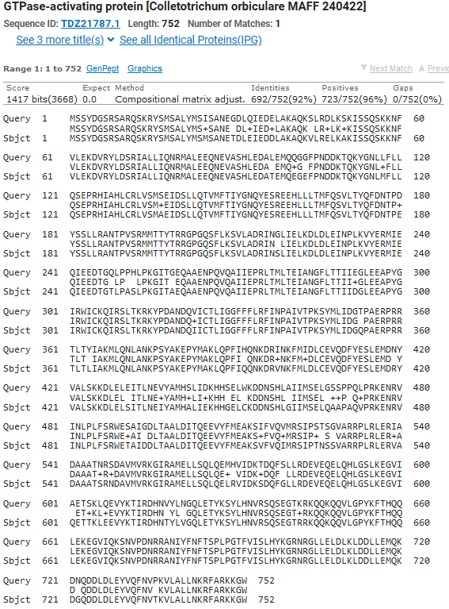
